# Girding the loins? Direct evidence of the use of a medieval parchment birthing girdle from biomolecular analysis

**DOI:** 10.1101/2020.10.21.348698

**Authors:** Sarah Fiddyment, Natalie J. Goodison, Elma Brenner, Stefania Signorello, Kierri Price, Matthew J. Collins

## Abstract

In this paper we describe a dry non-invasive extraction method to detect palaeoproteomic evidence from stained manuscripts. The manuscript analysed in this study is a medieval parchment birth girdle (Wellcome Collection Western MS. 632) made in England and thought to be used by pregnant women while giving birth. Using a dry non-invasive sampling method we were able to extract both human and non-human peptides from the stains, including evidence for the use of honey, cereals, ovicaprine milk and legumes. In addition, a large number of human peptides were detected on the birth roll, many of which are found in cervico-vaginal fluid. This suggests that the birth roll was actively used during childbirth. This study is the first to extract and analyse non-collagenous peptides from a parchment document using a dry non-invasive sampling method and demonstrates the potential of this type of analysis for stained manuscripts, providing direct biomolecular evidence for active use.

## Introduction

### History of the birth roll

Even today childbearing can be a highly perilous time for both mother and child [1–3], but in medieval Europe the risks were considerable. In eleventh-century Norwich, skeletal material from a more deprived area reveals the average age of death for women was thirty-three and the infant mortality rate over sixty percent [4]. In the latter Middle Ages, even though more women survived into old age as the standard of living increased considerably and childbirth was not the main cause of death for aristocrats [5], even the healthiest of women in childbearing had good reason to fear a protracted confinement, permanent injury, if not death [4]. The high death-toll for women, at what we would today consider an early age, reflects complications caused by postpartum infection, uterine prolapse, or other complications, and clearly indicates the act of giving birth was fraught with danger. Pre-Reformation English devotion indeed encompassed many feminine appeals for safe delivery, which included invoking St Mary (the mother of Jesus), St Anne (the mother of St Mary), and St Margaret, who was once swallowed by a demonic dragon and burst forth -- a sign taken in the Middle Ages of release from the prison of pregnancy [4]. But on the whole, we have very little surviving first-hand evidence from medieval women themselves about either the treatment or the complications of their own bodies [6,7]. The medieval Church offered a bevy of relics or talismans in various forms that were believed to aid in bringing about a safe pregnancy and delivery. A supplicant might gaze upon, touch, kiss, wear, recite, venerate, or even ingest this divine protection [8]. Midwives deployed parchment amulets, precious stones and plant-based remedies during childbirth [9]; the list of items that the church lent out to pregnant women is extensive. When Thomas Cromwell ordered the abbeys to be raided in what would come to be known as the Dissolution of the Monasteries, the raids were particularly vicious in targeting centres relating specifically to the veneration of female saints, much of which focused on childbearing and pregnancy [4]. The list of items siezed included St Moodwyn’s staff, which was lent out to women travailing, because *‘it was good for them to hold*’ [10]. Others include the smoke of St Mary, or even her breast milk [11]. However, the most oft-recited item that a monastery lent out to its wealthier parishioners was a birthing girdle.

These girdles seem to have taken a fair number of material forms. The girdle of Our Lady from Bruton was made of red silk [12]. At Coverham, the girdle of Mary Nevell and good for women lying in, was made of iron [13]. It seems that abbeys, such as Westminster, loaned out these girdles to women for a sum, such as the six shillings and eightpence paid “to a monke that brought our Lady gyrdelle to the Quene”, noted in Elizabeth of York’s Privy Purse Expenses, December 1502 [14]. Girdles had several functions. They were to be worn and unloosed before marriages, and a similar ‘girding and unloosening’ occurred during pregnancy: girdles may have literally supported the extra pregnancy weight and then been loosened six weeks prior to confinement [15].

However, it seems the birth girdle, especially birthing rolls such as MS. 632, was often talismanic, with ritual functions that spanned religious devotion and magic. Here, these artefacts formed part of a broader cultural landscape in which both women and men appealed to divine and supernatural forces for assistance in the face of ill health or danger. Devotional objects, such as relics or statues of saints, were venerated and touched in order to harness their beneficial power. Religious gestures like these sat alongside magical practices, most obviously the use of charms, formulae of which the efficacy was derived from the power of words [8]. Women’s health, encompassing not just pregnancy and birth but also menstruation and the various health issues shared with men (digestive complaints, problems with the eyes and ears, pestilential illness, etc.) was often addressed in the arena of oral culture, where religious and magical rituals were prominent. Miracle accounts are a rich source of information about behaviours that may otherwise be undocumented, and include references to the use of garments or belts associated with saints to assist women in labour. An account from the 1170s of a healing miracle of St Thomas Becket describes how Alditha of Worth, after a labour lasting three days, was finally able to deliver her infant when encircled by a stole that St Thomas had blessed [16]. Here, as in other contexts, emphasis is placed on the encircling role of the artefact, binding and protecting the maternal body.

Although this devotional and magical context is essential to understanding the cultural place of birth girdles, their use is also attested in medical texts, with two mentions in such texts of girding the loins of women in childbearing. *The Trotula* (a well-known medieval manual on women’s health), states that the skin of a snake should be hung about the abdomen. If a woman was having difficulty with a child coming out, ‘let the woman be girded with a snake’s skin from which the snake has emerged’ [17]. *The Sickness of Women*, another medical text, also recommends a birth girdle. For the grievances of women travailing in childbirth, one remedy recommends, ‘*and lete guyrden hir with a guyrdel of an hertis skynne*’ (and let her be girded with a girdle of hart’s [deer] skin) [18]. These girdles, of silk, iron, parchment, or snake skin, do appear to have been used by some pregnant women during the Middle Ages, especially by wealthy women, and were on the list of devotional items that were taken away during the Dissolution.

Perhaps because of the fervor with which the birthing talismans were destroyed by Reformers, or perhaps because of the delicate nature of the organic fibers that some other girdles were made from, few of these survive. The rolls that do survive, at least the ones that are known to us, are rolls mainly made from parchment, such as MS. 632 (Figure 1), although some printed on paper are also known (Harley MS 5919, held at the British Library) [19]. In addition to MS. 632, there are seven other English girdles known to us in the British Library and other collections, and at least two French girdles also held by Wellcome Collection.

**Figure 1.**
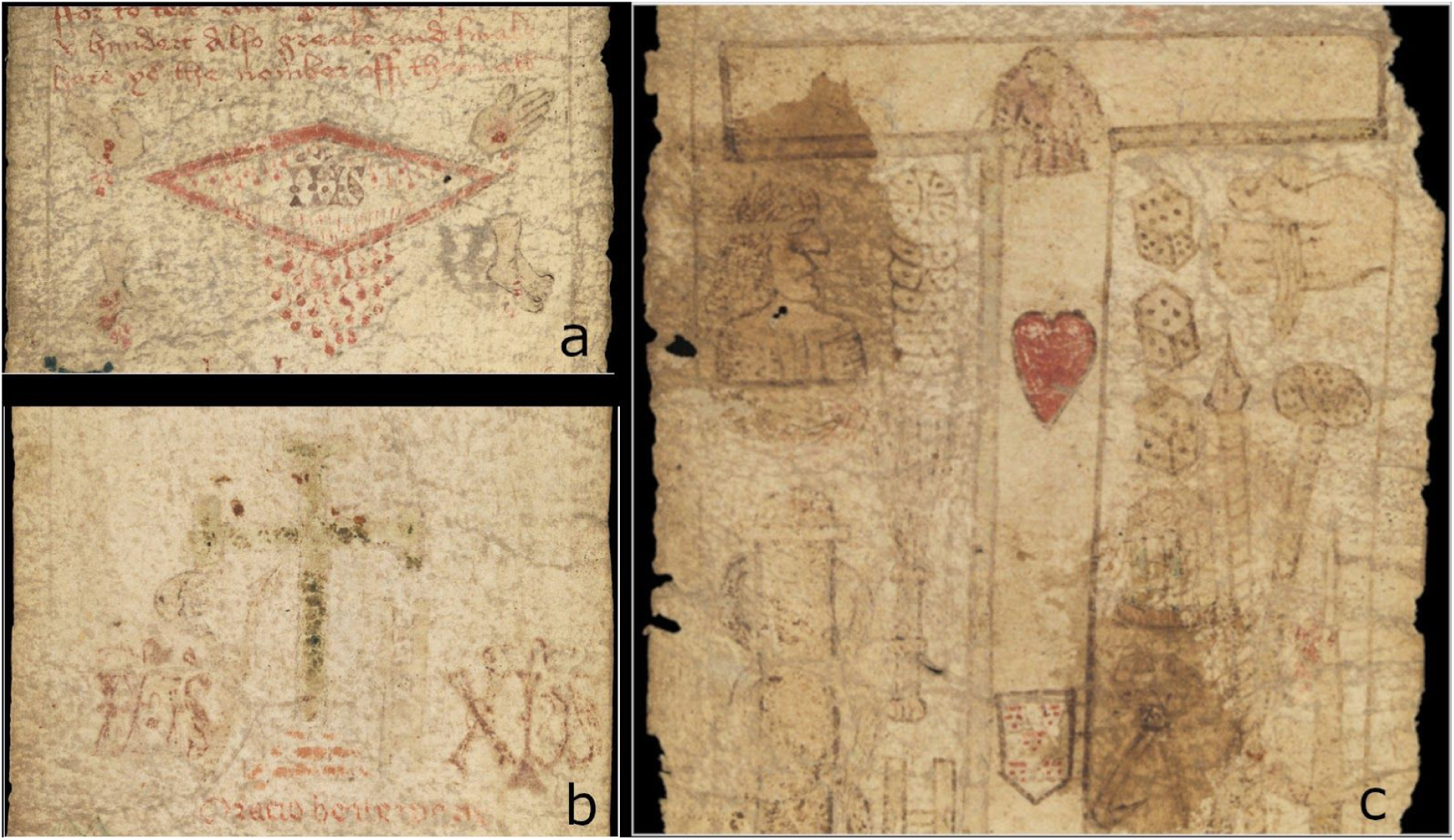
Details taken from MS. 632. **a**) MS. 632: the dripping side-wound. **b**) MS. 632: rubbed away green cross or crucifix. **c**) MS. 632: Tau cross with red heart and shield. Images courtesy of Wellcome Collection.

However, what makes MS. 632 largely unique among the remaining parchment rolls, is that it has obvious signs of actual use as a birth girdle. Several of its texts and crudely drawn images are associated with childbirth. It makes special invocation to Saints Quiricus and Julitta, mother and son martyrs, who in English birth roll traditions are invoked for aid specifically in childbirth [19]. The prayers in many of the rolls, MS. 632 included, are diverse --warding against many kinds of troubles-- and due to its various ‘catch-all’ nature of inclusion, the roll may have been used by men as well as women to guard against danger or hardship [20,21]. However, MS. 632 also crucially contains one final invocation specifically for women: ‘*And yf a woman travell wyth chylde gyrdes thys mesure abowte hyr wombe and she shall be delyvyrs wythowte parelle and the chylde shall have crystendome and the mother puryfycatyon*’ [And if a woman travailing with child girds this measure about her womb, she shall be delivered safely without peril and the child shall be christened and the mother purified]. Furthermore, the roll includes several invocations to St Mary, a saint who was understood to provide much assistance to women in pregnancy and childbirth [16]. One of these references invokes the tradition of the letter to Pope Leo, in which an angel delivered a letter to the Pope from the Virgin herself, transferring particular rites of protection to ‘*who so beryth ths mesure uppon hym*’ (the ‘mesure’ was thought to be the height of the virgin).

The writing on the manuscript appropriates the precise numbers, invocations, and repetitions found in magical talismans and enchantments into a Christian context. It includes the twelve names of the apostles, the names of the three magi, the repetition of precise numbers (the exact number of the drops of Christ’s blood), and the use of sacred holy names, both in invocation and in abbreviation (such as the holy initials of Christ). Such features indicate that the highly formulaic rituals blend magic rituals with religious protection [8].

MS. 632’s severe abrasions implies that it was often touched or kissed; and its narrow width (330.0 x 10.0 cm) suggests that it was intended to imitate an actual metal or cloth girdle that could be wrapped around a woman’s body, with the strategic placement of particular prayers against her womb [19]. The folds in the parchment of MS. 632 certainly attest to such a type of arrangement and the length of the manuscript would make this possible. If the roll was wrapped around the waist, draped down the backside, lifted up through the legs and draped back over the woman’s abdomen toward her chest, this would fit the cross-like shape suggested by Gwara, Morse, Jones and Olsan, and also demonstrate how it would be plausible to completely gird the woman with the roll during the act of labour (Figure 2).

**Figure 2.**
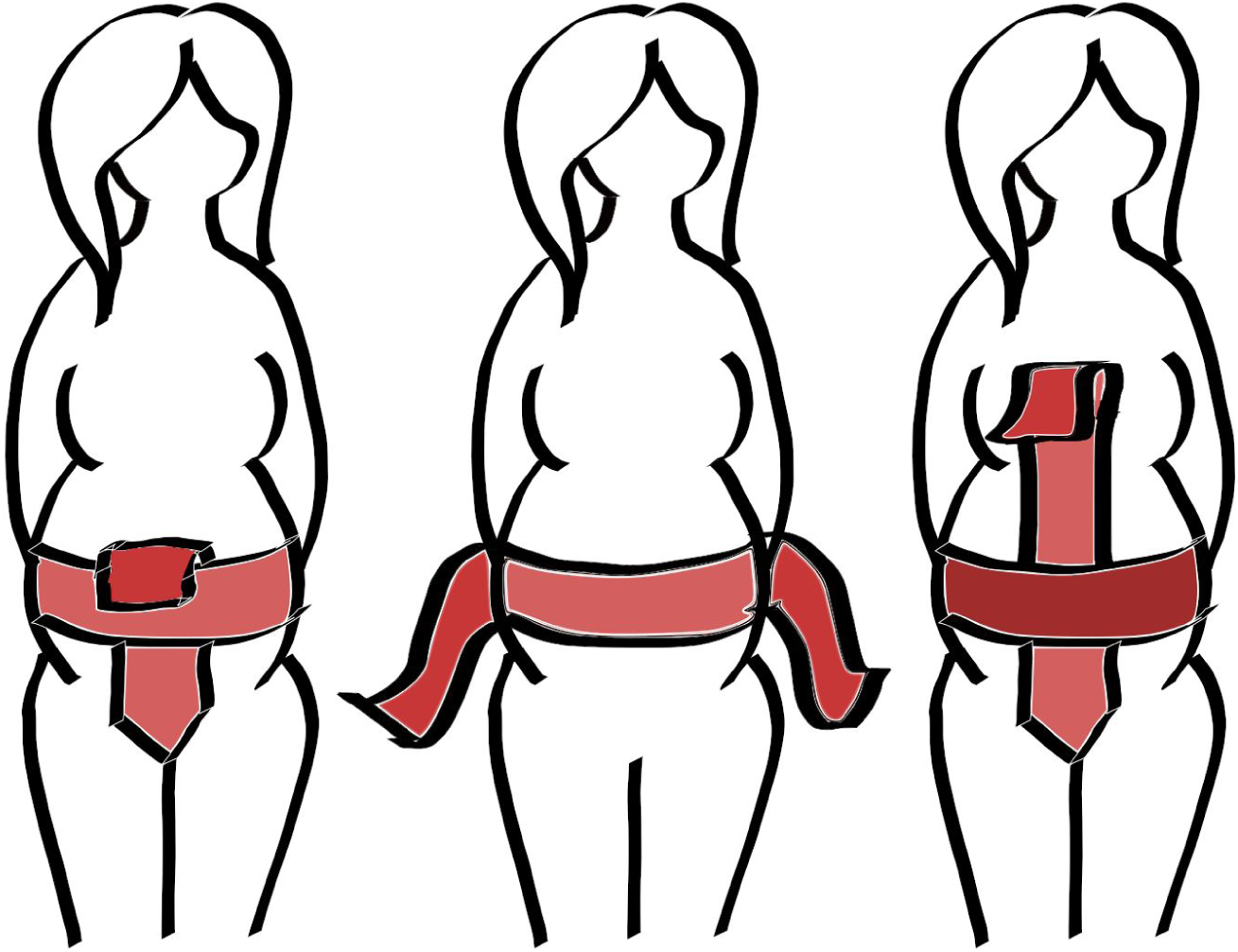
Possible methods of tying the birth girdle when used pre and during labour.

The overall effect of the manuscript is that it has been very heavily worn, often thought through use, with the significant abrasions on the surface of the parchment making large sections of the text illegible. This most probably indicates that it has frequently been used as a birth girdle [22]. In fact, such abrasions and wear has even called for MS. 632 to be tested to see if its use in childbirth can be scientifically verified. Lea Olsan writes: ‘Were they damaged during repeated use in the events surrounding labour and delivery, when the roll, according to its instructions, was laid over the womb of the woman to ease the delivery? There are a few reddish marks that could be blood stains where the roll is very worn, but laboratory work will be required to make certain’ [23]. In this work we take up Olsan’s challenge and attempt to provide identification on the stains on these samples.

### Recent advances in palaeoproteomics

Recent advances in palaeoproteomic techniques have allowed a much more in-depth examination of biomolecular evidence on different substrates. Proteomic analysis of bone [24–26], tooth enamel [27,28] and shell [29] have proved the survival of a diverse collection of proteins in addition to the primary structural protein. This holds great promise for phylogenetic studies where genetic analysis is not possible due to poor preservation of DNA [27,30]. By contrast proteins appear to have a much longer survival rate (well over one million years) [29] giving the tantalising possibility of pushing the limits of biomolecular analysis much deeper into time.

Proteomic analysis has also been successfully carried out on more recent objects but with equally interesting outcomes. Initial proteomic studies of cultural heritage objects has always required taking physical destructive samples [31]. In addition to the obvious obstacle of obtaining destructive samples, there is an inherent bias to the predominant protein present (in the case of parchment that protein is collagen) and consequently the signal of lower concentration proteins is drowned out, in most cases to such a degree that they are undetectable.

### Non-invasive techniques

The use of non-invasive EVA diskettes on numerous paper documents [32–34] and even mummified skin [35] has provided interesting details about the use of these objects, and in some cases possible evidence of pathogens has been detected. It has recently been used for the first time on parchment [36], however due to the nature of the sampling (requiring the damp diskette to be in contact with the parchment) it may limit its widespread adoption. Here we report the first proteomic analysis of a historic parchment document using a dry non-invasive sampling technique (eZooMS) developed previously for species identification of parchment. The technique has now been expanded to analyse the broader set of proteins present on the surface of the document which can provide information about the history and use of this object.

## Results and discussion

### eZooMS

All eight eraser samples were analysed via MALDI-TOF analysis for species identification. In all cases the species of parchment was determined to be sheep (Figure 3). We can conclude therefore that all four separate pieces of parchment that make up the birth roll are all made from sheep parchment. Previous studies have shown a predominance of sheep parchment in English legal documents [37] however, in this case the reason for using sheepskin may be for more practical reasons. Given that sheepskin is more affordable than calfskin, and more readily available in England than goat, it would seem an obvious choice. More importantly, sheep skin is much thinner than calfskin, lending itself to make documents that need to be folded or manipulated, such as in the case of the birth girdle.

**Figure 3.**
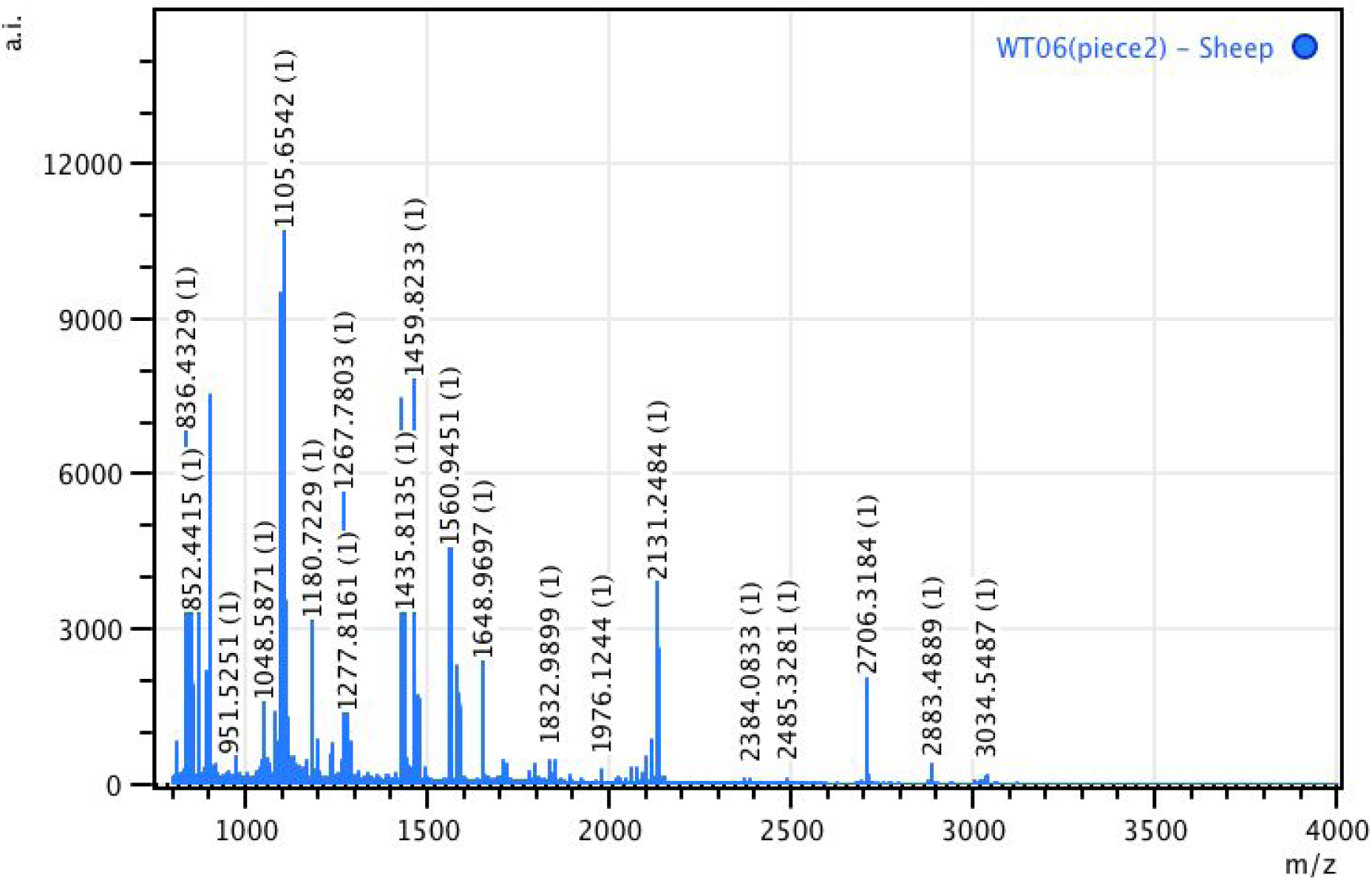
Example of MALDI-TOF spectra from one of the birth girdle samples (WT06) identified as sheep parchment.

### LC-MS/MS

A list of proteins found in the birth roll samples can be found in Supplementary info Table 1. Of the 273 proteins found in the birth roll samples, 201 were found exclusively in these samples. Only proteins that were found exclusively in the birth roll samples are discussed in the results. Differential expression therefore was not taken into account in this study (merely presence/absence), but there is the potential for these studies to be conducted at a later stage.

**Table 1.** shows the proportion of proteins (all and exclusive) present in the different samples.

| Sample | All proteins | Exclusive prot | Human proteins | Human excl prot | All human proteins found in CVF | Excl proteins found in CVF |
| --- | --- | --- | --- | --- | --- | --- |
| WT | 273 | 201 | 100 | 54 | 50 | 25 |
| Blank (Paper blank) | 6 | 1 | 3 | 0 | 3 | 0 |
| Control (parchment control) | 73 | 2 | 48 | 2 | 25 | 0 |

**Figure 4.**
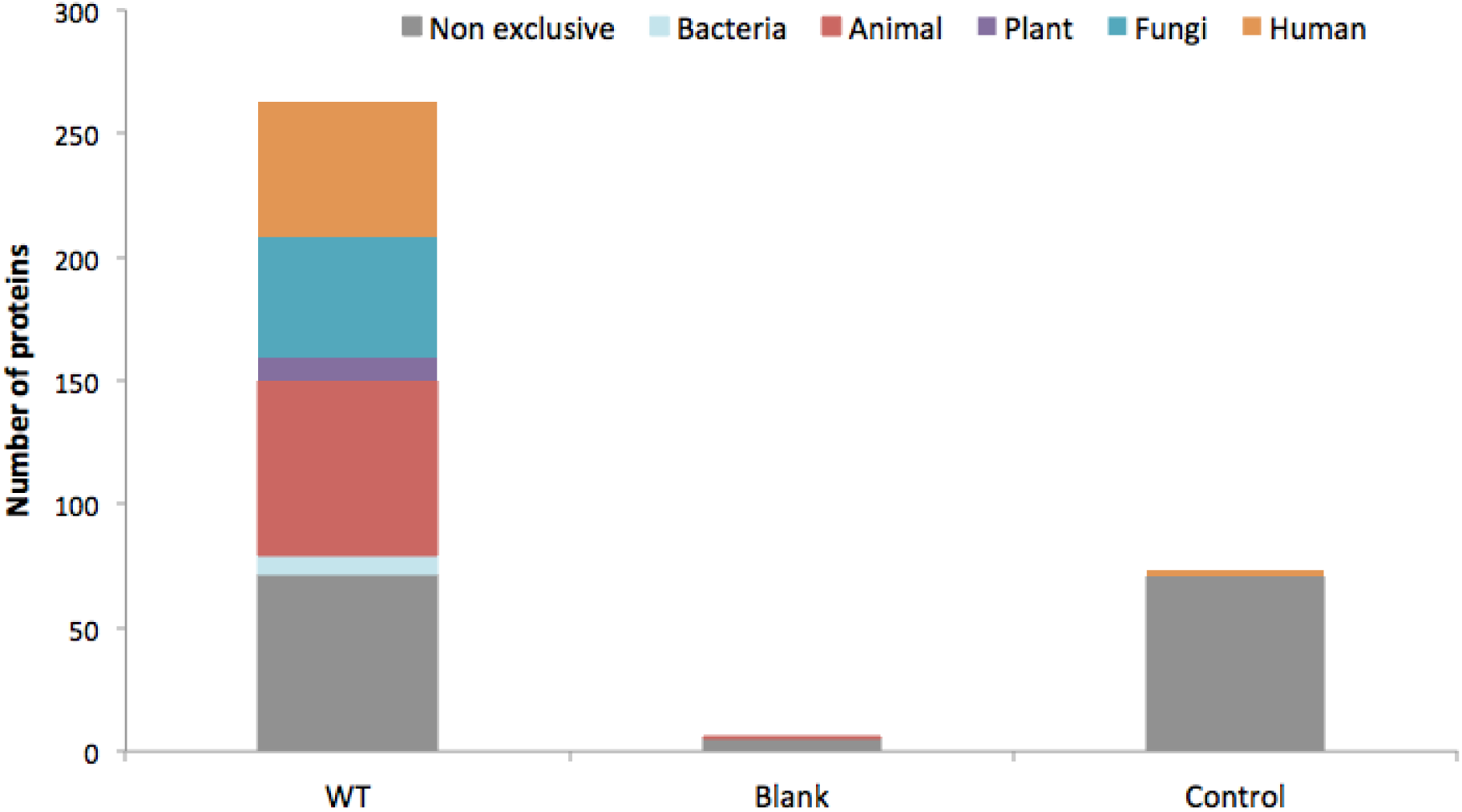
Shared proteins and exclusive proteins by category found on the three groups of samples analysed: Birth girdle (WT), Blank (paper) and Control (18th C parchment).

### Non-human proteins

‘*Take one scruple of opium poppy, one scrupe of goose fat, four scruples each of wax and honey, one ounce of oil, the whites of two eggs, and the milk of a woman. Let these be mixed together and inserted by means of a pessary*’, (Trotula, p. 89).

We were able to detect the presence of a number of animal derived proteins including honey (royal jelly protein) in WT04, ovicapra milk derived products in WT07 (casein), egg white (ovalbumin) and egg yolk (although two peptides also found in the control). We were also able to detect the presence of leguminous plants (broad beans and possibly garden pea), as well as cereals.

**Figure 5.**
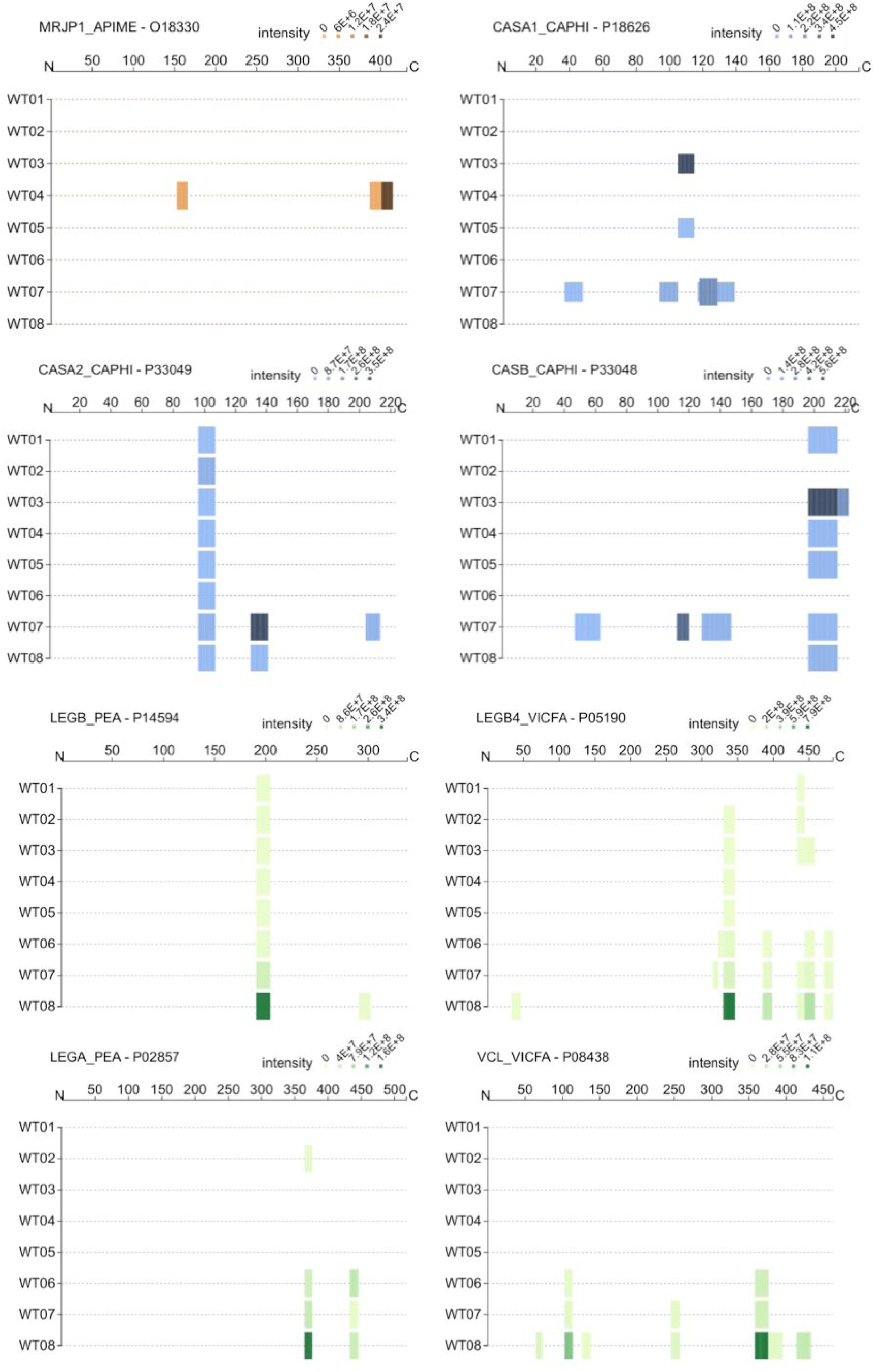
Plots showing all the peptides present for the main non-human proteins detected in the birth girdle samples using Peptigram [46]. Each coloured band represents one peptide sequence and the intensity of colour reflects the peptide intensity detected.

### Honey

One of the birth girdle samples (WT04) presented three peptides (IMNANVNELILNTR, LLTFDLTTSQLLK, MVNNDFNFDDVNFR) identified exclusively to the Major royal jelly protein 1 (MRJP1, ID:O18330) from *Apis mellifera* (Honeybee).

Honey has been used throughout history for medicinal purposes, both for its natural antiseptic properties as well as it’s sweetness to make mixtures more palatable. There is evidence in the Trotula that it was also used to remedy problems in childbirth including for movement of the womb from its place. For lesions of the womb, ptisan and honey were combined and inserted into the womb as a pessary [17]. Honey again is used on poultices to tie up the womb after it has come out after birth. Herbs (garden cres, laurel berries, frankincense, and cinnamon) and pulverized and mixed with honey, placed on the loins and tied with a band in a ball, and inserted into the vagina [17].

### Egg

Vitilogenin-2, a protein found in egg yolk was detected in two of the eight birth roll samples. However, two (different) peptides were also observed in our control parchment and therefore we cannot exclude some form of contamination. However, two peptides from Vitilogenin-1 were exclusively found in the birth roll samples, making a stronger case for the presence of egg yolk.

There is evidence of the use of egg yolk during childbirth, and the complications surrounding women’s health, in medieval treatises, with one mention in the *Trotula* recommending that to avoid a difficult birth the mother should eat easily digestible foods for the last three months of pregnancy, including yolks of egg [17]. In addition there is later 17^th^ C evidence, but that may reflect earlier medieval knowledge, recommending that birth mothers should eat certains foods to keep their strength up during labour, including a poached egg yolk [38–40]. In another, egg yolks are used as lubricant to facilitate delivery in the case of a difficult labour [38,40–42].

The presence of egg white was detected in all eight birth girdle samples but not in the blank or control. Egg white is often used to treat the surface of the parchment [43], so in the case of the birth roll we cannot conclude categorically that it’s presence is due to medicinal purposes as it could equally be explained by virtue of the preparation of the parchment. In addition, chicken lysozyme is detected in many of the same samples (lysozyme is also found in egg white).

There is also evidence for the use of egg whites found in medieval recipes for women’s health. One reference in the *Trotula* regards the use of egg whites for ulcers of the womb, thought to be occasioned by miscarraige. To mitigate the pain, the text indicates the one should apply the ‘juice of a deadly nightshade, great plantain with rose oil, and white of egg with woman’s milk and with purslane juice and lettuce,’[17]. Another recipe regarding ‘*excessive heat of the womb*’ uses two of the items found on the Wellcome birth girdle. ‘*Excessive heat of the womb*’ draws on the Hippocratic of bodily humours: women were normally cold and moist, but pregnancy could upset this balance. If the womb was thought to be too hot, this remedy was applied: ‘Take one scruple of opium poppy, one scrupe of goose fat, four scruples each of wax and honey, one ounce of oil, the whites of two eggs, and the milk of a woman. Let these be mixed together and inserted by means of a pessary’, [17]. This uses honey and egg whites, both found in the analysis.

### Milk

Ovicaprine milk was detected in WT07: alpha-casein 1 (five specific peptides), alpha-casein 2 (two peptides) and beta-casein (five peptides). Milk has been described as a surface treatment of parchment [43,44], however, we found no evidence of milk on the control parchment which would seem to indicate that the presence is due more to the nature and use of the birth girdle. It is interesting to note, unlike beta lactoglobulin (BLG), the milk protein most commonly reported in dental calculus, that the only milk protein detected was casein. In separated milk, BLG is enriched in the whey, while casein is concentrated in curds, and therefore also present in cheese and other milk derived products, with any one of these being the possible origin.

There is evidence in the Trotula suggesting ‘...writing symbols on cheese or butter and giving them to the mother to eat, in order to hasten a delivery’[17,45], which could offer a possible explanation for the presence of casein on the birth girdle.

### Plants

#### Leguminous plants

We detected seven peptides from vicilin from *Vicia faba* (broad bean) in the three of the birth girdle samples. In addition we also detected the presence of peptides from two other proteins, Legumin A (two peptides) B (two peptides), and Legumin type B (eight peptides) although the specificity to organism can not be differentiated between broad bean, garden peas and common vetch.

Leguminous plants were frequently used in medieval recipes, including women’s health. For example, *The Trotula* uses beans for lesions of the womb, to instigate the flowing of breast milk, and for curing certain pregnancy cravings. Some of these recipes include the mixing the bean with wheat and barley--which too are detected in this manuscript. *The Sickness of Women* uses bean meal as a dietary remedy for dropsy and for swollen legs during pregnancy; beans and peas were used to increase fertility in both men and women. Considering legumes were used in medieval recipes for childbearing, its presence in MS. 632 may be explained through medieval remedies surrounding childbirth. The legumes’ presence may even suggest what types of ingredients were used in England as remedies to problems surrounding childbearing.

#### Cereals

We were also able to detect the presence of cereals in the birth girdle samples, although many of the peptides were not specific enough to determine the exact species, the closest matches included wheat, barley and spelt. There is the possibility that their presence could be explained by the use of a conservation treatment using wheat or flour paste. However, it would be unlikely that they used all of the cereals and we have yet to find documentary evidence that any conservation of this kind has been undertaken. Cereals can also be used as a surface treatment in parchment production [43] and could therefore explain their presence.

Yet there is ample evidence of cereals, including wheat and barley, used for treatments surrounding women’s health and childbirth. Barley was a particularly versatile ingredient, used in remedies for excessive menstrual blood flow, menstrual pain, lesions of the womb, vaginal wind, the expedition of a delivery, and as a contraceptive. For difficult deliveries, the *Trotula* instructs a woman to be bathed in water in which barley, mallow, fenugreek, and linseed have been cooked. In another instance in the *Trotula*, if a woman was badly torn and did not wish to give birth again, she must put barley or grains of caper spurge into her afterbirth, the amount of grain for the number of years she wished to remain barren. Wheat flour was used to stimulate a maternal milk-flow, to treat lesions of the womb, and to determine male or female cause of infertility [17]. While the presence of cereals on the birth girdle may indicate parchment preservation, it may also indicate their active use in treatment surrounding childbirth thereby affirming evidence that such recipes as present in the *Trotula* were widely used.

### Human proteins

A total of 54 human proteins were found exclusively in the birth roll samples, compared to two in the control sample. These proteins were then compared to the proteome from cervico-vaginal fluid (CVF) published by Muytjens et al [47].

A higher number of proteins from the birth roll samples are detected in CVF than from any other of the sample groups. This can provide a further plausible indication that the roll was indeed actively used during childbirth (Figure 6).

**Figure 6.**
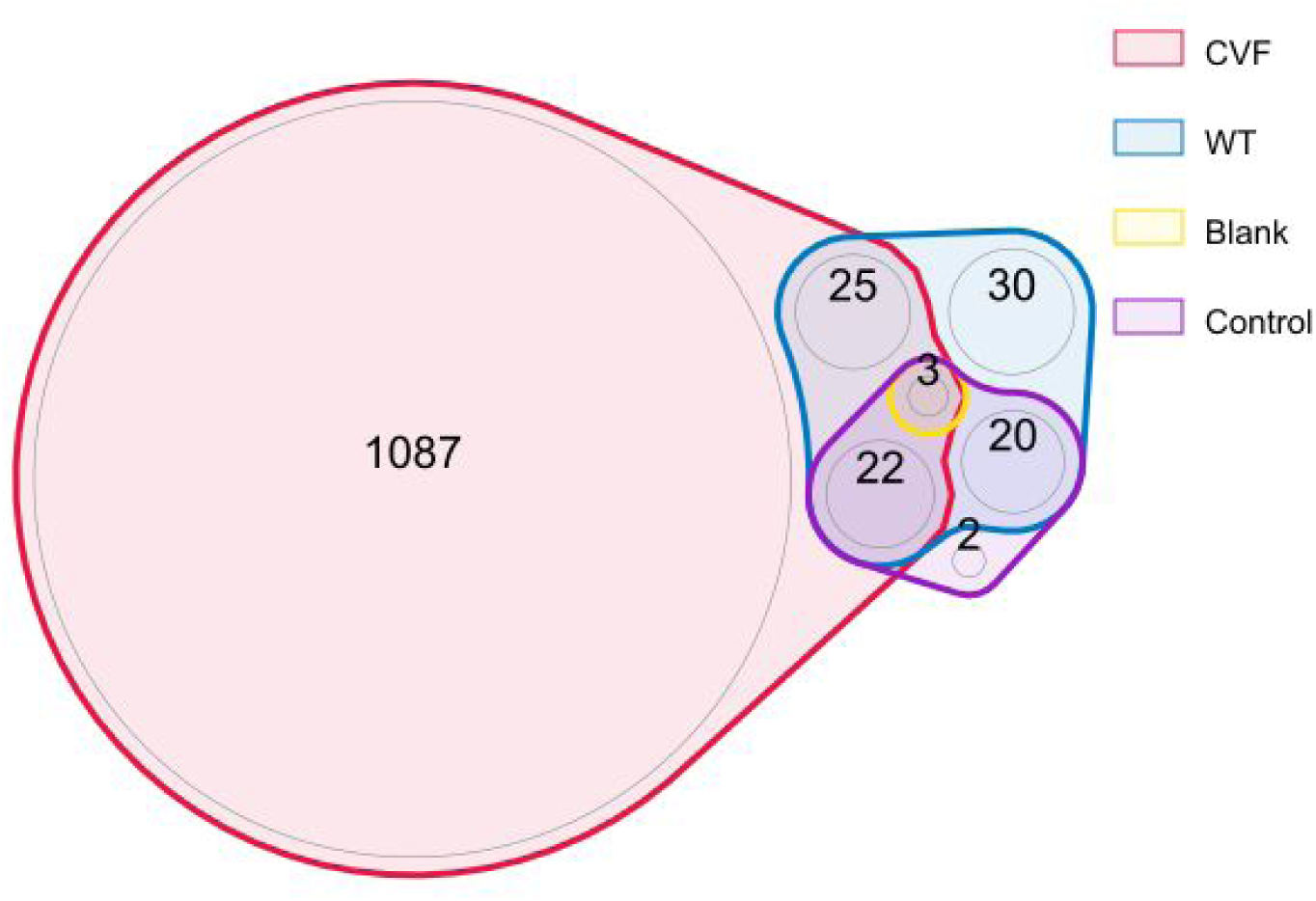
Venn diagram of human proteins present in the different samples compared to proteins found in cervico-vaginal fluid (CVF). WT (birth girdle) samples have the largest number of human proteins, and the largest number of exclusive proteins found in CVF, compared to the blank and control.

As detailed above, the assumption about the method of use of birth rolls is that they were worn during childbirth by the mother, possibly tied around the waist like a girdle, acting as a talisman or good luck charm for what has always been a significantly dangerous and potentially fatal experience for women. If indeed the roll was used in this way, it would be logical to expect to find traces of bodily fluids and medical remedies present on the roll.

In addition, we were also able to detect nine peptides belonging to major urinary proteins from *Mus musculus* (mouse) which would indicate that during the history of the object, at some point it has been stored in place where mice have had access to it and left this evidence. However, we cannot say when this happened. In addition, we recorded a large number of peptides belonging to different strains of *Aspergillus*, something that has also been noted by other researchers looking at parchment [36]. *Aspergillus* seems to often be found colonising parchment samples, but to what degree this may be a conservation concern requires further research.

Remembering the words of Lea Olsan cited in the introduction, obtaining direct evidence of use of these birth girdles is almost impossible save for the idea that direct biomolecular data could be extracted and identified in a laboratory setting. We have now shown that with current non-invasive methodologies it is possible to obtain direct evidence of use that provides clear data to support the historical scholarship of birth girdles.

## Conclusion

Non-invasive proteomic analysis of manuscripts can provide a deeper understanding of the history and use of the object. Surface sampling preferentially extracts substances that have been deposited on the document and avoids the inherent bias from predominant proteins when using physical destructive samples. Proteomic analysis has the advantage over genetic analysis in that it’s tissue specific providing information not only on species but materials used. We have demonstrated its applicability with the example of this medieval birth roll providing proteomic evidence of its possible active use during childbirth. We have also proved what was long suspected of MS. 632, that its badly worn state attests to its use during childbirth. The use of the honey, legumes, eggs, and even milk-products are also common remedies in childbirth and this study lends further support to medieval medical treatises, such as those recorded in obstetric manuals, were practices that were actively employed. The active use of MS. 632 in childbirth also shows that women were using highly formulaic rituals that blended the numerical precision and incantation of magic with religious protection. The potential of proteomic analysis applied to the vast corpus of parchment documents represents a huge new avenue of exploration for the burgeoning field of biocodicology.

## Materials and Methods

### Samples

Wellcome MS. 632 is a long and thin roll, dating to the late fifteenth century or early 1500s, measuring 332 cm x 10 cm. It is made of four strips of parchment stitched together. The roll contains:1-the three nails of the crucifixion; 2-the crucifix with a heart and a shield; 3-a circle surrounding the Holy monogramme, ‘IHS’ (the first three letters of the name ‘Jesus’ in Greek--iota (i), eta (h), sigma (s)); 4-the hands and feet of Christ (i.e. the five wounds of Christ) dripping with blood; 5-a very rubbed crucifix, faded; 6-a diamond-shaped figure; very badly rubbed, possibly Christ standing [20–22]. A unique point relating to this girdle is that it has text on both the face and dorse of the roll, which is unusual.

### eZooMS

Eight non-invasive eraser samples were taken from Wellcome Library MS. 632, in both stained and unstained areas. At least one sample came from each of the four different pieces of parchment that make up the roll in order to determine the species of animal used to make the parchment in each of the cases. Samples were taken using PVC erasers as described in Fiddyment et al. 2015. Samples were extracted using conventional eZooMS methodology [37] and initially analysed using MALDI-TOF MS. Eraser crumbs collected from rubbing the eraser on a blank sheet of paper acted as a blank, and rubbings taken from an 18th C Scottish legal document acted as a control.

### LC-MS/MS

The remaining peptides left from the eZooMS analysis were dried down in an evaporator and sent for analysis to the University of Copenhagen (Globe Institute & Centre for Protein Research) where they were analysed via LC-MS/MS following the Copenhagen protocol from Demarchi et al. [29].

### Data analysis

Data was analysed using the open source software MaxQuant [48] (version 1.6.10.43) with tandem mass spectra searched against the Uniprot_SwissProt database (downloaded 23/01/2020) including in the search all common contaminants (cRAP). Mass tolerance was set to ±5 ppm on the precursor and 0.05 Da on the fragments. The thresholds for peptide and protein identification were set as follows: protein false discovery rate (FDR) = 0.01, protein score and peptides matches ≥ 1. Subsequent analysis was then carried out using the Proteus R package (https://github.com/bartongroup/Proteus) [49]. Only proteins with ≥ 2 peptides were considered during the results and analysis.

The mass spectrometry proteomics data have been deposited to the ProteomeXchange Consortium via the PRIDE partner repository [50] with the dataset identifier PXD022054.

## Acknowledgements

The authors would like to thank Meaghan Mackie at The Globe Institute, University of Copenhagen for her help with running the LCMS/MS samples and her advice on data preparation. The authors would also like to thank the Institute of Medieval and Early Modern Studies for facilitating a portion of this collaboration.

## Authors’ contribution

SS & SF came up with the initial question and SS undertook the non-destructive sampling. SF extracted and analysed all the samples. SF and MJC interpreted all the data. EB, NG and KP conducted all the historical contextualisation. All authors gave final approval for publication and agree to be held accountable for the work performed therein.

